# Iron limitation by transferrin promotes simultaneous cheating of pyoverdine and exoprotease in *Pseudomonas aeruginosa*

**DOI:** 10.1101/2020.06.21.163022

**Authors:** Oswaldo Tostado-Islas, Alberto Mendoza-Ortiz, Gabriel Ramírez-García, Isamu Daniel Cabrera-Takane, Daniel Loarca, Caleb Pérez-González, Ricardo Jasso-Chavez, J Guillermo Jiménez-Cortés, Yuki Hoshiko, Toshinari Maeda, Adrian Cazares, Rodolfo García-Contreras

**Author notes:** These authors contributed equally (Oswaldo Tostado-Islas, Alberto Mendoza-Ortiz, Gabriel Ramírez-García, Isamu Daniel Cabrera-Takane, Daniel Loarca, Caleb Pérez-González). Correspondence: R García-Contreras, Departamento de Microbiología y Parasitología, Facultad de Medicina, Universidad Nacional Autónoma de México, 04510, Mexico City, Mexico. E mail, Adrian Cazares, EMBL’s European Bioinformatics Institute (EMBL-EBI), Wellcome Genome Campus, Hinxton, Cambridge, United Kingdom. E mail.

## Abstract

*Pseudomonas aeruginosa* is the main bacterial model to study cooperative behaviors, since it yields exoproducts such as exoproteases and siderophores that act as public goods and can be exploited by selfish non-producers that behave as social cheaters. Non-producers of the siderophore pyoverdine are typically isolated in media with low free iron, mainly casamino acids medium supplemented with transferrin. Nevertheless, using a protein as the iron chelator could additionally select mutants unable to produce exoproteases that degrade the transferrin to facilitate iron release. Here, we investigated the dynamics of pyoverdine and exoprotease production in media in which iron was limited by using either transferrin or a cation chelating resin. Our experiments show that concomitant loss of pyoverdine and exoprotease production readily develops in media with transferrin whereas only lack of pyoverdine emerges in medium treated with the resin. Genomic characterization of the exoprotease- and pyoverdine-less mutants revealed large deletions (13 to 33 Kb) including Quorum Sensing (*lasR, rsal and lasI*) and flagellar genes. Complementation experiments, PCR and motility tests confirmed the deletions. Our work shows that using transferrin as an iron chelator imposes simultaneous selective pressure for the loss of pyoverdine and exoprotease production. The unintended effect of transferrin observed in our experiment settings can help revisiting or informing the design of similar studies.

## Introduction

*Pseudomonas aeruginosa* is a major bacterial model to study cooperative behaviors since it produces a variety of exoproducts, such as siderophores, which are considered public goods [1]. These molecules are important iron scavenging agents, essential for iron acquisition in iron-limited media; however, its production is susceptible to exploitation by social cheaters, since siderophores are released to the environment and can benefit both producer and non-producer individuals. Most of the published works aiming to study siderophore cheating in *P. aeruginosa* have used casamino acids (CAA) medium supplemented with 100 μg/mL of human apo-transferrin (CAA TF) in order to chelate iron and make iron acquisition and growth dependent on siderophore production[2–7]. This medium was originally proposed by Griffin and coworkers in 2004. In 2017 Harrison and collaborators compiled a list of published experiments on siderophore cheating in *P. aeruginosa* to that date, covering 23 articles and 36 experiments, of which 34 were performed in CAA TF [2]. These works include studies on the long-term coevolution between cheaters (PAO1 *pvdD*) and cooperators (PAO1 wild-type) pyoverdine cheating and cheating resistance in isolates from soil and ponds and the dynamics of exoprotease and pyoverdine non-producers in media with protein as sole carbon source and under iron limitation[8].

More recently, the use of the CAA TF medium has been reported in research addressing the role of transposable phages in the evolution of social strategies in iron-limiting medium [9], the effect of cheating in diversity of pyoverdine variants and the effect of nutrients availability in the dynamics between wild-type and non-siderophore-producing mutants [4].

The CAA TF medium has been used assuming it creates conditions specifically suitable for the selection of non-siderophore producers, nevertheless, it is well known that quorum sensing (QS)-dependent exoproteases such as elastase LasB and alkaline protease AprA greatly enhance the rate of iron acquisition through pyoverdine both *in vitro* and *in vivo* [10, 11]. Hence, likely during infections, iron acquisition may depend on both siderophore production and exoproteases activity. Exoproteases are also public goods, and therefore susceptible of exploitation by non-producers [12–16].

In most studies of pyoverdine dynamics, iron in the medium has been limited by the action of apo-transferrin, a protein susceptible to degradation by *P. aeruginosa* QS-regulated exoproteases; hence, we hypothesized iron acquisition in such conditions is mediated by two types of public goods: siderophores (pyoverdine) and QS-regulated exoproteases that facilitate iron release from the degraded transferrin. Consequently, the production of both pyoverdine and exoproteases would be susceptible to be cheated by non-producers, favoring the selection of these mutants. Here we show that limiting iron availability by transferrin (TF) promotes strong selection of exoprotease non-producers in addition to pyoverdine-less mutants. Our experiments identified different phenotypes emerging from continuous growth in CAA TF medium in terms of pyoverdine and exoprotease production, but double cheaters of these exoproducts were prevalent at the end of the experiments. Whole genome sequencing and analysis of three mutant types allowed the detection of small sequence variants associated to loss of pyoverdine production but also uncovered extensive genome deletions in the exoprotease non-producers. The deletions included QS genes, essential to produce exoproteases, as well as genes encoding diverse functions such as flagellum components. Hence, the use of transferrin as iron chelator in our experiment settings drove the emergence of exoprotease-less individuals through large genomic rearrangements affecting phenotypes other than siderophore production, an unintended effect that might have influenced similar studies.

## Methods

### Strains media and growth conditions

The strains used in this work and their characteristics are summarized in Table 1.

Bacteria was cultured in CAA medium, containing per liter: 5 g casamino acids, 1.18 g K2HPO_4_.3H_2_O, 0.25 g MgSO_4_.7H_2_O and 25 mM HEPES buffer, CAA TF medium (CAA medium supplemented with 20 mM NaHCO3 and 100 ug/mL of human apo-transferrin), CAA chelex medium (CAA treated with chelex 100 resin according the manufacturer instructions (5 g of resin per 100 mL of medium). Minimal succinate medium (MSM per liter: 6 g K_2_HPO_4_, 3 g KH_2_PO_4_, 1 g (NH_4_)_2_SO_4_, 0.2 g MgSO_4_·7H_2_O, and 4 g succinic acid), and MSM TF (MSM supplemented with 20 mM NaHCO_3_ and 100 ug/mL of human apo-transferrin) Final pH was 7-.0.

Bacteria were cultured in Erlenmeyer flasks of 50 mL o capacity filled with 5 mL of medium, at 37°C with 200 rpm orbital shaking, during 18 h for growth experiments and 24 h for continuous sub-culturing, each new culture was initiated with enough bacteria to reach an O.D. 600 nm of 0.05.

### Isolation of pyoverdine and exoprotease less individuals

Single colonies were obtained by streaking each subculture made in a pair day in LB plates that were further incubated at 37 °C for 18 h, after that 20-50 colonies were passed to King A plates and were incubated under the same conditions, those that were green and showed fluorescence under UV light were considered pyoverdine producers. Also, colonies were passed to M9 minimal medium plates with casaminoacids 0.025 % (w/V) and 0.5 % (w/V) casein and were incubated under the same conditions; those that produced a clear halo (indicating casein degradation) were considered exoprotease producers. Some single colonies exhibiting different phenotypes were picked, grown in liquid CAA medium and stored in 16 % glycerol at −70 °C for further experiments.

### Bacterial competitions

In order to evaluate the cheating capacity of some of the isolated colonies that were non-producers of pyoverdine and or non-producers of exoprotease, they were incubated in CAA, CAA TF and M9 Casein medium in co-culture with the PA14 wild-type strain at 15-20% as an initial amount and incubated during 24 h, samples were taken just after inoculation and at 24 h of growth and colonies were isolated for phenotypic determination (pyoverdine and exoprotease production) in order to calculate the initial and final proportions.

### Iron determination

The content of Fe in growth media was determined by digesting 5 mL of each one with 5 mL of concentrated nitric acid with overnight incubation at 90°C. The content of metals was determined by atomic absorbance spectrometry (ASS) by using a Varian apparatus as reported elsewhere.

### Pyoverdine, exoprotease and pyocyanin measurements

Pyoverdine from the bacterial supernatants was measured fluorometrically, using a Perkin Elmer Victor Nivo plate reader, exciting at 405 nm and recording at 510 nm.

The exoprotease activity found in the supernatants was determined by measuring hydrolysis of Azo-casein (SIGMA, St Louis Mo, USA) as described in [17], using also a Perkin Elmer Victor Nivo plate reader determining absorbance at 415 nm.

Pyocyanin was extracted from the supernatants using chloroform, re extracted with HCL 0.2 N and determined spectrophotometrically as previously described [18].

### Swimming motility assays

Swimming plates were made with 0.25% (W/V) agar Luria broth (LB). After solidification, plates were briefly dried at room temperature. A spot inoculated with 2.5 μL aliquots taken directly from overnight LB cultures. Swim plates were incubated at 37°C for 18-20 h. An image of the swimming plate was captured with a digital camera.

### Whole genome sequencing and variant calling analysis

Genomic DNA used for sequencing was extracted from overnight cultures. Library synthesis was performed using the Nextera XT DNA Sample Prep Kit (Illumina, San Diego CA, USA) according to the manufacturer’s instructions. TruSeq HT adapters (Illumina, San Diego CA, USA) were used to barcode the libraries, and each library was sequenced using an Illumina MiSeq 300 bp paired-end instrument. Trimmomatic v.0.39 [19] was used to process the raw fastq files for adapters and quality trimming with the following settings: “LEADING:3, TRAILING:3, SLIDINGWINDOW:4:23 MINLEN:35”.

Single nucleotide polymorphisms (SNPs) and small indels were detected with the Snippy v3.2 pipeline (https://github.com/tseemann/snippy) by aligning the sequencing reads to the genome of the reference strain UCBPP-PA14 (Accession: NC_008463.1) with BWA-MEM (Li, H. Aligning sequence reads, clone sequences and assembly contigs with BWA-MEM, calling variants with FreeBayes [20] and annotating them with snpEffm [21]. The coverage metrics of the genes *lasR* and *pvdS* were determined with samtools v.1.7 [22]. Deletions flanking *lasR* were inferred from coverage distribution data obtained with BEDTools [23] as the longest stretch of gene coding regions featuring zero reads. De novo assembly of the genomes was carried out using SPAdes v.3.10.1 [24] with default settings. Visualization of coverage distribution and inspection of the joining regions was performed with Artermis [25]. The genome assemblies are publicly available from the BioProject PRJNA540579.

### Complementation of the strains

The construction pUCP20-*lasR* was donated by Professor Gloria Soberón Chavez from the Institute of Biomedical Research at UNAM, for the pUCP20-*pvdS* construction, *pvdS* gene was cloned by amplifying it from the PA14 genomic DNA, standard methods were used to isolate chromosomal DNA [26]. Restriction DNA enzymes and T4 DNA ligase were purchased from England Biolabs (NEB, Ipswich MA, USA) or Invitrogen. Plasmid used for cloning the sigma factor *pvdS* (was purified using an E.Z.N.A Plasmid mini kit I, (Q-spin) (Omega Bio-Tek). DNA was amplified with the appropriate oligonucleotides using Phusion High-Fidelity DNA polymerase (Thermo scientific, Sweden) according to the manufacturer’s recommendations. The resulted *pvdS* obtained by PCR was purified by E.Z.N.A. Cycle pure kit (Omega Bio-tek). pUCP20-*pvdS* plasmid carries a 742pb fragment that includes the coding region of *pvdS* (564 pb), 151 pb upstream and 27pb downstream. For this, *pvdS* was amplified by PCR using the oligonucleotides *pvdS*-Fw: 5’GCTCTAGAGCCAGCATGCGGACCATTCAC3’ and *pvdS-Rv:* 5’GCGAATTCACCGGCGCTGAGGAATGC3’ The product was then cloned into pUCP20 as an XbaI-EcoRI fragment, the resultant plasmid pUCP20-*pvdS*, was inserted by transformation into *P. aeruginosa.* Complementation of the *lasR*-less phenotype was done in the presence and absence of 5 μM 3-Oxo-C_12_-HSL (Sigma) and pyoverdine, caseinolytic activity and pyocyanin was determined as explained previously.

### Transformation of *Pseudomonas aeruginosa* by CaCl_2_

Briefly, *P. aeruginosa* was grown overnight in Luria Broth (LB) medium with orbital shaking (200 rpm) at 37°C. 250 μL of the cells culture were harvested by centrifugation at 15000 x g 1 min, washed in same volume of cold sterile Milli-Q water. Then the cell suspension was centrifuged at 15000 x g 1 min and the supernatant was discarded. The cell pellet was suspended in 250 μL of 0.1 mM CaCl_2_ and incubated for 20 min at 4°C. The cells were centrifuged at 15000 x g 1 min. The supernatant was retired and the cell pellet suspended in 100 μl of 0.1 mM CaCl_2_ whit 100 ng of the pUCP20-*pvdS* construction or the empty plasmid. The suspension was maintained at 4°C for 1 h. Cells were heat pulsed by partially immersing in water at 42°C for 2 min. After the heat step the cells suspension was incubated at 4°C for 2 min. 500 μL of LB medium was added to the cells and then incubated at 37°C for 1 h. Transformant colonies were selected on King-B plates [27] containing 300 μg/mL carbenicillin.

### PCRs for the confirmation of deletions

The following PCR reactions were done in order to confirm the presence of genomic deletions: amplification of a fragment of 742 pb bp of the gene *pvdS* as positive control, using the primers *pvdS*-Fw 5’GCTCTAGAGCCAGCATGCGGACCATTCAC 3’ and pvdS-Rv 5’GCGAATTCACCGGCGCTGAGGAATGC 3’, to confirm the integrity of the chromosomal DNA isolated from the wild-type *P. aeruginosa* as well as the mutants CH1, CH2 and CH3, b) fragment of 720 bp of the gene *lasR*, using the primers F- lasR 5’ATGGCCTTGGTTGACGGTT 3’ and R-*lasR* 5’GCAAGATCAGAGAGTAATAAGACCCA 3’. All the PCR reactions was done like previously described.

### Statistical analysis

Results shown are the average of at least 3 independent experiments ± SD or SEM, data was analyzed using the IBM SPSS Statistics software and the statistical tests: Mann-Whitney, Kruskall-Wallis, and Student T-test were applied to different data sets as specified in the figure legends, differences were considered significant when P values were lower than 0.05.

## Results

### Selection of siderophore and exoprotease non-producers in the presence of apo-transferrin

Colonies of the *P. aeruginosa* strain PA14 were grown and serially transferred in CAA medium supplemented with apo-transferrin (CAA TF) for 14 serial passages and periodically tested for pyoverdine and exoproteases production. Siderophore-less mutants were identified from passage number 6 (~24 generations). Figure 1A, shows that in parallel to the selection of pyoverdine-less mutants, there was a strong selection of exoprotease non-producers. A screening of single colonies of the two phenotypes revealed that most of them exhibit concomitant loss of pyoverdine and exoproteases production (Figure S1).

**Figure 1.**
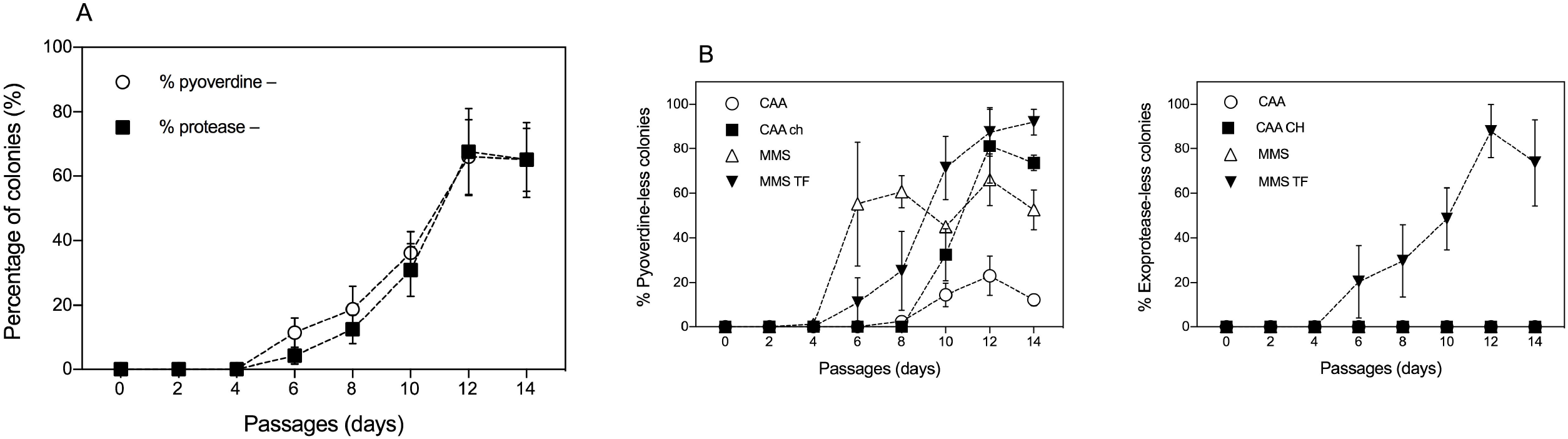
A) Percentage of pyoverdine and exoprotease non-producers of the PA14 strain PA14 identified during sequential subculture in CAA medium supplemented with transferrin (CAA TF). Results are the average of 8 independent experiments ± SEM. No significant differences between the percentages of non-pyoverdine and non-protease producers were found using a Mann-Whitney U test. B) Percentage of pyoverdine (top) and exoproteases (bottom) non-producers of the strain PA14 identified during sequential subculture in CAA medium, CAA medium treated with chelex 100 (CAA ch), minimal succinate medium (MMS) and minimal succinate medium supplemented with transferrin (MMS TF). Results are the average of 3 independent experiments ± SEM.

To test if the presence of apo-transferrin was responsible for the selection of exoprotease-less mutants, a new set of experiments was done in the CAA medium using the chelex 100 resin as the iron-depletion factor. Non-treated CAA and minimal succinate medium (MSM) were also used as a control and as an alternative of iron-limited medium, respectively. Our results show that pyoverdine non-producers appeared from passage 6 (Figure 1A and B) and were selected upon passage 14 in all media; however, they were more abundant in the CAA medium treated with chelex, and in MMS, than in the non-treated CAA medium (Figure 1B), as expected from the lower iron availability in those media (13.1 ± 3.25, 7.12 ± 1.8, and 66.3 ± 24.7 μM for CAA chelex, MMS, and CAA respectively). We did not find exoprotease non-producers in the screening of bacteria grown in media lacking apo-transferrin (Figure 1B). In contrast, adding apo-transferrin to the MSM promoted the selection of exoprotease-less mutants in parallel to pyoverdine non-producers (Figure 1B), thus confirming that the presence of apo-transferrin imposes a selective pressure for the loss of exoprotease production. The addition of TF to the CAA medium also promoted the emergence of exoprotease- and pyoverdine-less individuals in sequential cultures of the reference strain PA01 (Figure S2), thus suggesting that this selection may not be exclusive to the strain PA14.

### Characterization of pyoverdine and exoprotease non-producer individuals

Our experiments demonstrated that the CAA TF medium selects the loss of both pyoverdine and exopreotease production, hence, four different types of individuals are expected to occur in the population: wild-type (wt, pyoverdine and exoprotease producers), pyoverdine cheaters (pyoverdine non-producer, exoprotease producers), exoprotease cheaters (pyoverdine producer, exoprotease non-producers), and double cheaters (pyoverdine and exoprotease non-producers). The presence of the four types of individuals was verified in the experiment in CAA TF medium and their frequency from passage 6 to 14 was recorded. We found that double cheaters were the predominant cheater type from passage 8 and that they reached around 90% of the population in the passage 14 (Figure S1), hence implying that these individuals have a greater fitness than those losing a single trait.

Next, we characterized a set of individual colonies by measuring production of pyoverdine and exoproteases (caseinolytic activity), and growth in different media. The colonies selected were representative of the three different cheater phenotypes: one colony identified as pyoverdine and exoprotease non-producer (CH1), other identified as pyoverdine producer but exoprotease non-producer (CH3), and the last colony identified as pyoverdine non-producer but exoprotease producer (CH2). We found that the growth of the three isolates was similar to PA14 wild-type in CAA medium; however, the pyoverdine non-producers (CH1 and CH2) grew less than the wild-type strain in iron-limited media (CAA TF and CAA chelex) (Figure S3). Likewise, exoprotease non-producers (CH1 and CH3) grew less than the wild-type strain in the medium containing casein as sole carbon source (Figure S3). As expected, the levels of fluorescence in the pyoverdine non-producers CH1 and CH2 were considerably lower than in the CH3 isolate (Figure 2 A). The caseinolytic activity assay also confirmed the reduction in exoprotease activity in CH1 and CH3 but not in the CH2 strain (Figure 2B). Competition experiments between PA14 wild-type and two different CH strains showed that CH1 can cheat both pyoverdine and exoprotease since its frequency in co-cultures increased after 24 hours in CAA TF and M9 Casein media but not in non-treated CAA medium (Figure 3A), whilst CH2 increased its frequency only in an iron-limiting condition (Figure 3B).

**Figure 2.**
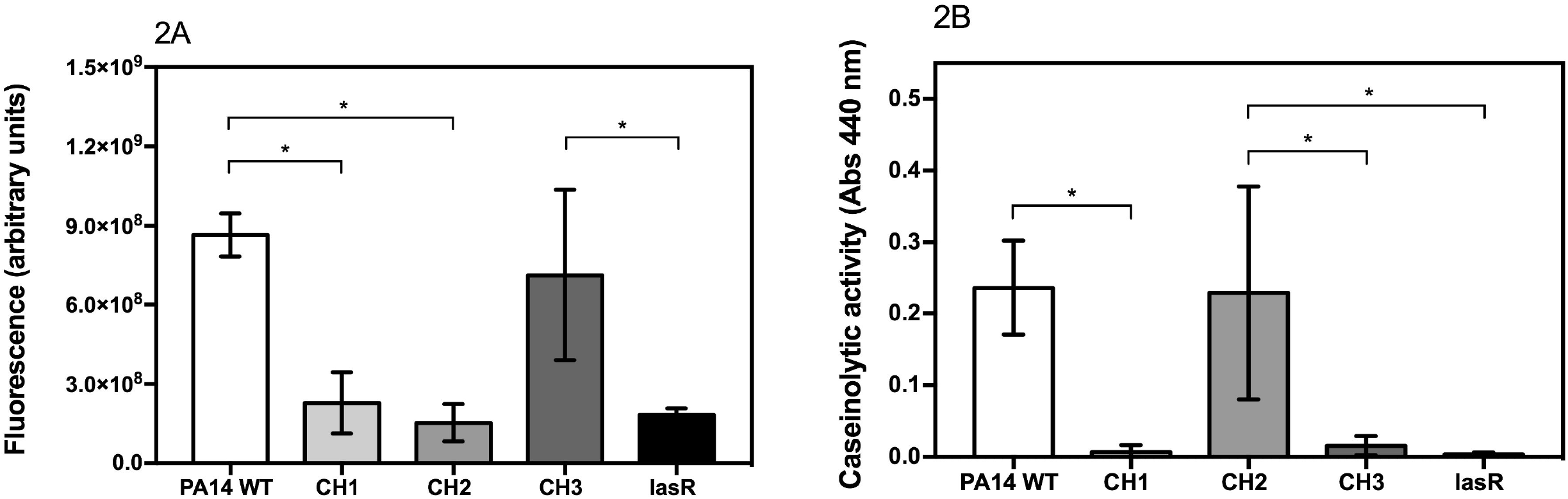
A) Pyoverdine production of the PA14 wild-type strain, 3 PA14-derived isolates from the evolution experiments (CH1, CH2, CH3), and a *lasR* deletion mutant. Plotted are the averages of at least 5 independent experiments ± SEM. Significant differences in production of pyoverdine between PA14 WT and CH1 (p=0.023), PA14 WT and CH2 (p=0.001), CH2 and CH3 (p=0.019), and PA14 WT and *lasR* (p=0.005) were found using a Krustall-Wallis test (p<0.001) with Bonferroni correction. B) Exoprotease production (caseinolitic activity) of the PA14 wild-type strain, 3 PA14-derived isolates from the evolution experiments (CH1, CH2, CH3), and a *lasR* deletion mutant. The values plotted correspond to the average of at least 5 independent experiments ± SEM. Significant differences in exoprotease production between PA14 WT and CH1 (p=0.005), PA14 WT and CH3 (p=0.002), CH1 and CH2 (p=0.015), CH2 and *lasR* (p=0.007), and PA14 WT and *lasR* (p=0.002) were found using a Krustall-Wallis test (p<0.001) with Bonferroni correction.

**Figure 3.**
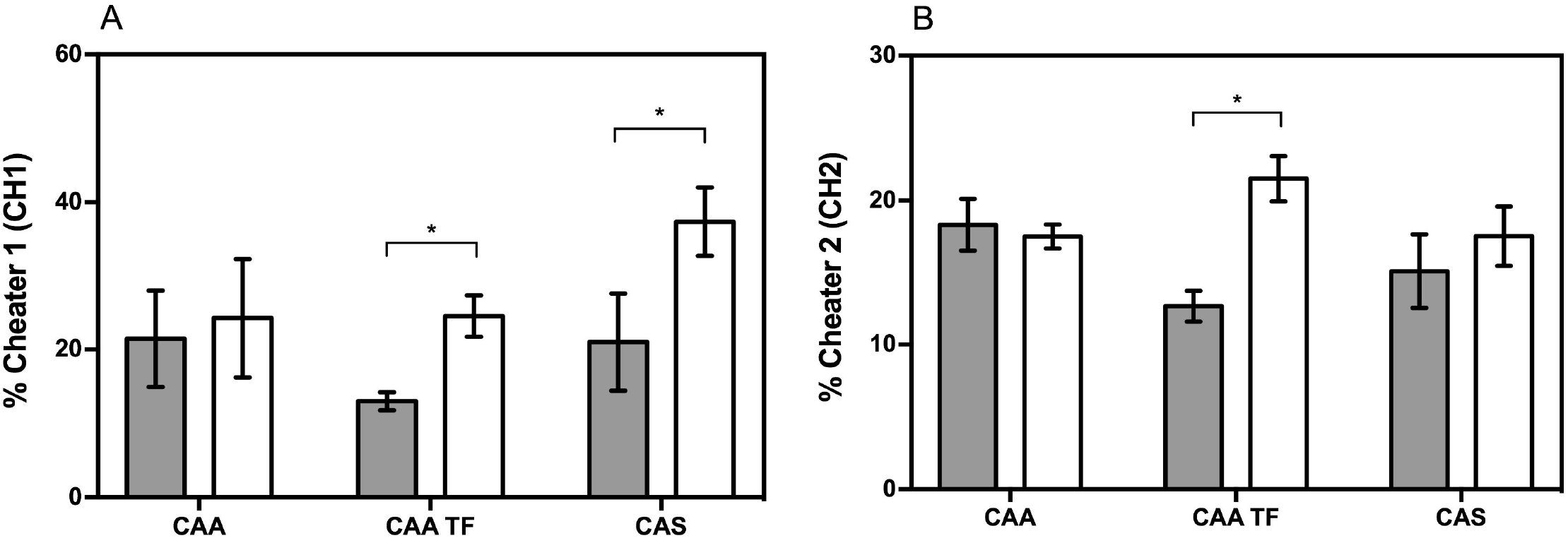
Competitions between the PA14 wild-type strain and CH1 (A) or CH3 (B) in CAA medium, CAA medium supplemented with transferrin (CAA TF), and caseinate medium (CAS). The barplots show the average ± SD of the initial (grey bars) and after 24 h (White bars) proportion of CH1 in the competition experiments. The experiments were performed in triplicate. The differences between the initial and after 24 h proportions of CH1 in CAA TF and CAS media are significant according to T-test, P >0.05. Difference between proportions of CH3 are significant in CAA TF medium.

### Whole genome sequencing

To get insights into the genetic basis of the loss of exoprotease and pyoverdine production, we performed whole genome sequencing of the PA14 wild-type strain used in this work, the three CH isolates, and an additional clone with wild-type phenotype isolated from the CAA-TF experiments (WT1). A variant calling analysis targeting small indels and nucleotide polymorphisms identified few mutations occurring in the CH isolates but not in the wild-type and WT1 strains (Supplementary Table 1), nevertheless, their association with the observed phenotypes was not entirely clear. Unlike previous reports [3, 9, 14, 15, 28], we did not detect mutations in the *pvdS* and *lasR* genes which are commonly associated with loss of pyoverdine and exoprotease production. The pyoverdine non-producer isolates CH1 and CH2, however, featured a frameshift mutation in *pvdA* and one mutation in the intergenic region upstream the *pvdS* gene, respectively. To assess whether low sequencing coverage hampered the identification of mutations in the regions encoding LasR and PvdS, we determined the coverage metrics for these genes in all the isolates (Supplementary Table 1). The *pvdS* sequence was covered by hundreds of reads in all the genomes with comparable RPKM values. In contrast, very few or no reads mapping *lasR* were detected in the genomes of the exoprotease non-producers CH1 and CH3. Detailed examination of the *lasR*-flanking regions in both genomes revealed several genes with zero reads coverage, indicative of large deletions (Figure 4). The coverage assessment of the regions predicted the loss of ~33 and 13 kb in the CH1 and CH3 genomes, respectively (Figure 4). The putative deletions included flagellar genes, components of an RND multidrug efflux pump, and the quorum sensing genes *lasI, rsaL* and *lasR* (entirely or partially) (Supplementary Table 1). Additionally, the deletion in CH1 covered some transporters, regulators, and a larger number of flagellar genes, among others.

**Figure 4.**
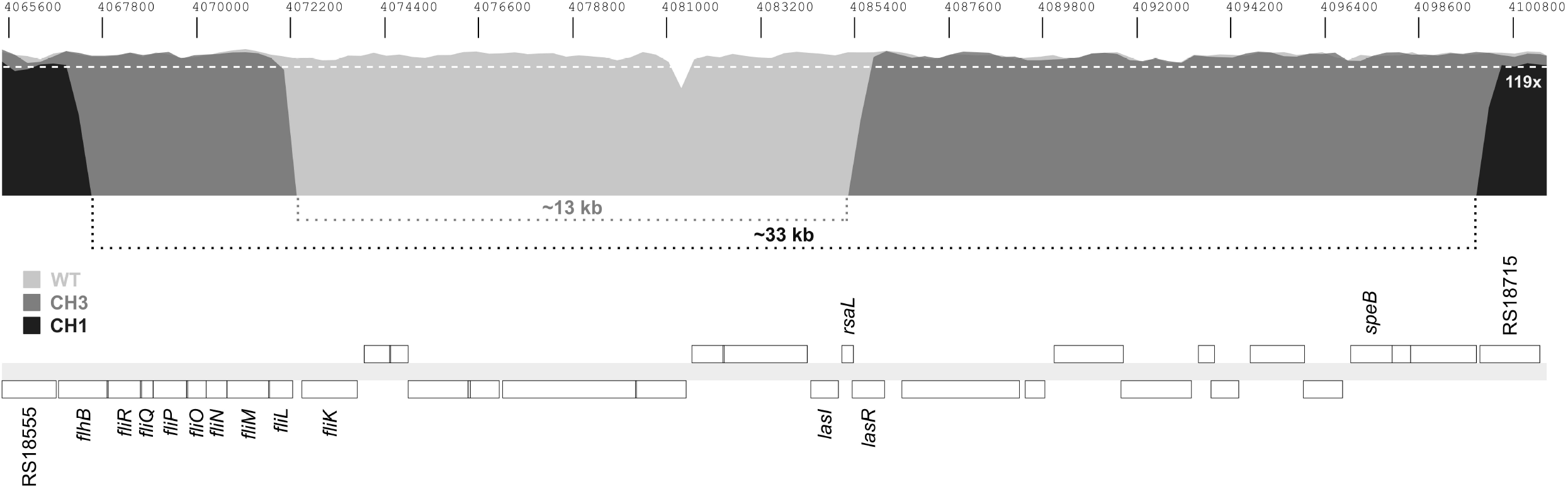
Multigene deletions on the genomes of the isolates CH1 and CH2. The figure shows the coverage distribution of sequencing reads mapping a selected region of the PA14 genome (NC_008463). Coordinates of the selected region are displayed above the sequencing coverage plot. Coverage by reads from the CH1, CH2, or wild-type (WT) genomes is color coded and indicated in the figure. A white dotted line in the plot indicates a reference coverage value. The position and approximate length of regions featuring zero mapped reads from the CH1 and CH2 genomes, indicative of deletions, is marked by dotted lines below the coverage plot. A gene map of the PA14 selected region is shown at the bottom of the figure with genes represented by white boxes. Names or locus tags of genes of interest are indicated above or below their corresponding boxes.

### Confirmation of genome deletions and motility tests

We designed primers targeting the genes *lasR* and *pvdS* to validate the deletions identified in the CH1 and CH3 isolates. The fragment corresponding to the amplification of *lasR* was detected in the strains CH2 and wild type but not in CH1 and CH3 (Figure S4). Conversely, *pvdS* was amplified in all the strains, as anticipated from our genome analysis. Since deletions in CH1 and CH3 include several flagellar genes, we further assessed the functional impact of this genomic loss by evaluating the swimming motility of the isolates. Neither CH1 nor CH3 exhibited motility, in contrast to the wild type and CH2 strains which lack the genome deletion (Figure S5).

Next, we assembled the CH1 and CH3 genomes to identify the newly generated joining regions and better define the structure of the deletions. Contigs containing junctions between the regions flanking the deletions were recognized in the two genomes (Figure 5). Detailed examination of the junctions revealed that in CH1 the deletion removed 33.2 kb of genome sequence including 30 coding regions and different portions of the genes *flhB* and RS18715 where the chromosome breaks occurred. In CH3, the intergenic region between *fliL* and *fliK*, and the gene *lasR*, correspond to the chromosome breakpoints of the 13.5 kb deletion that removed 11 genes and 157 of the 240 codons in *lasR.* In CH3 the breaks rejoin merging *the flhB* and RS18715 coding sequences whereas in CH1 the junction likely leads to loss of the *lasR* ORF coding potential.

**Figure 5.**
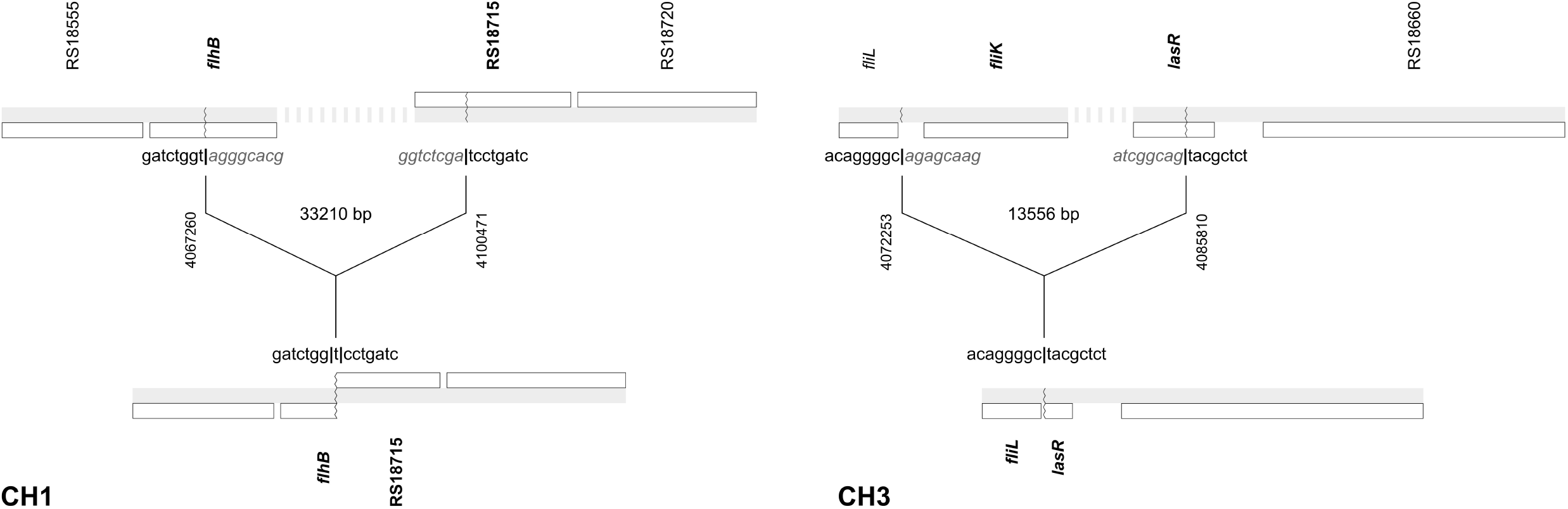
Junctions between regions flanking multigene deletions in the CH1 and CH3 genomes. Diagrams in the figure illustrate the chromosome breaks (top) and joining regions (bottom) of deletions identified in the genomes of the CH1 (left) and CH3 (right) isolates. Gene maps in the diagrams are draw to scale according to the annotations and coordinates of the PA14 genome (NC_008463). Genes are represented by white boxes. Gene names or locus tags are shown next to their corresponding boxes. The deletion breakpoints in the chromosome are depicted as zigzag lines in the maps and the genes affected by them are marked in bold typeface. The nucleotide sequences flanking the breakpoints prior deletion (top) and after junction (bottom), are presented in the figure. The position of the breakpoints prior deletion is indicated at the center of the diagrams along with the length of the deletion. The joining regions displayed in the figure were identified in contigs resulting from the assembly of the CH1 and CH3 genomes.

### Complementation of cheater phenotypes

Our genome sequence analysis revealed large deletions in the strains CH1 and CH3 encompassing the Quorum-sensing (QS) genes *lasI* and *lasR* (partially or entirely). Multiple studies on exoprotease cheating report that non-producers selected in medium containing protein as sole carbon source are commonly characterized as *lasR*-less mutants. Since the isolates carrying genome deletions lack *lasI*, we hypothesized that loss of exoprotease production could be complemented by an exogenous *lasR* only with the addition of the 3-Oxo-C_12_-HSL signal. Consistent with this notion, the transformation of CH3 with the construction pUCP20-*lasR* led to recover exoprotease production only in the presence of 3-Oxo-C_12_–HSL in the medium (Figure S4A). Pyocyanin production, also dependent on functional QS, was determined in the CH3 strain carrying pUCP20-*lasR*, with and without the 3-Oxo-C_12_-HSL signal. The results showed that CH3 is deficient in pyocyanin production, which can be complemented with pUCP20-*lasR* only when the signal is added (Figure S6A). Likewise, CH1 recovered exoprotease production in the presence of pUCP20-*lasR* and 3-Oxo-C_12_-HSL (Figure S6B). We also tested whether the CH2 isolate (pyoverdine non-producer, exoprotease producer) was defective due to lack of expression of the sigma factor pvdS, finding that complementation with an exogenous *pvdS* largely alleviates its deficit in pyoverdine production (Figure S6C).

## Discussion

Siderophore cheating has been extensively studied in the laboratory using *P. aeruginosa* as the main model since it represents an important and frequent behavior in communities of free-living bacteria [29]. Lack of pyoverdine production is also common, and increases with colonization time, in clinical *P. aeruginosa* isolates from cystic fibrosis patients that retain the ability to uptake it hence becoming potential cheaters. Similarly, *P. aeruginosa* QS-deficient mutants unable to produce exoproteases are commonly isolated from infections [30] in coexistence with QS-proficient isolates [31]. Loss of public good production and cheating seem to be a frequent phenomenon in *P. aeruginosa* and may contribute to attenuate its virulence during chronic infections; in fact, inoculating social cheaters in infections has been proposed as an strategy to attenuate host damage [32], supported by experimental evidence from animal infection models [33].

The strong selection for loss of exoprotease production upon cultivation with apo-transferrin presented here suggests that iron acquisition in such conditions not only depends on pyoverdine production, as initially assumed, but also relies on the activity of exoproteases that break down the transferrin, facilitating the release of the iron bound to it and enhancing the rate of iron acquisition through pyoverdine [10]. Since both siderophore and exoproteases are public goods, mutants that loss the expression of one or both traits, are benefited by the production of those factors by the cooperators, thus exploiting them.

Our genome analyses revealed that the loss of exoprotease production in the CH1 and CH3 mutants is not due to punctual mutations in *lasR* but rather to extensive genome deletions including the *lasR* and *lasI* genes; accordingly, complementation of exoprotease production in these mutants was only achieved by adding *lasR* in the presence of 3-Oxo-C_12_-HSL (Figure S4 AB). Moreover, there was a cluster of flagellar genes among the genes included in the deletions, consistent with the lack of swimming motility detected in the exoprotease-less individuals. In contrast with the loss of exoproteases in CH1 and CH3, loss of pyoverdine in CH2 was complemented by the addition of *pvdS*, in agreement with a mutation on the putative promoter region of this gene. Similar mutations affecting *pvdS* have also been found in other studies [3].

The lack of motility developed in some of our isolates is in line with previous findings reporting the inactivation of flagellar genes through punctual mutations in both pyoverdine non-producers and cooperators when the PA01 strain is serially cultured in CAA TF [3] and suggests that the lack of motility could be beneficial in such culture conditions. One possible explanation is that swimming does not confer an advantage during growth with shaking but the synthesis and function of the flagellum is energetically costly, therefore, losing it would allow the bacterium to save energy for other processes. Intriguingly, no deletion including *lasIR* genes was reported in the Kummerli et al. study [3]; we sought assessing the presence of deletions affecting the *lasIR* genes in the genome sequences presented in this work, however, these were not publicly available.

Other studies have found mutations in motility genes upon sub-culturing in CAA and CAA TF media [9]. Likewise, the archetypical strain to study siderophore cheating PAO6609 harbor mutations in motility genes, mutations in *lasI*, and a deletion of 6 kb in a region that includes genes involved with the synthesis of the pyoverdine side chain [2]; hence suggesting that losing QS and likely the ability to make exoproteases may be favorable for cheating in this mutant, and indicating that deletions can also be responsible for loss of pyoverdine production.

Experimental evolution studies investigating the development of resistance against the non-redox iron III mimetic gallium also involve the use of apo-transferrin to limit iron in CAA medium. We consider that selection for loss of exoprotease production represents a key variable likely overlooked in the experiments of these studies. In line with our findings, Bonchi and colleagues [34] reported a weak correlation between pyoverdine production and the ability of *P. aeruginosa* clinical strains to grow in human serum, nevertheless, a strong correlation between growth and exoprotease production was identified. The positive correlation between growth and proteolytic activity was due to degradation of the transferrin present in the serum leading to facilitation of iron release and uptake by pyoverdine, which translated into reduced susceptibility to the inhibitory effects of gallium [34].

Recent research has shown that coexistence between wild-type cells and pyoverdine (*pvdS* mutants) and exoprotease *lasR* mutants) non-producers is stable in low-iron conditions with casein as sole carbon source since the medium used in the experiments of this study contains apo-transferrin for achieving iron limitation[8], it would be interesting to test whether the same stability is observed when the iron availability is limited by methods other than the utilization of iron chelating proteins.

While others have shown that CAA TF do not necessarily resemble the growth conditions of *P. aeruginosa in vivo* here we show that strong selection for exoprotease-less mutants during sequential cultures in this medium may have also been overlooked, hence, we propose the utilization of other strategies for generating iron-limiting media. In our case, the use of iron-chelating resins removed 80% of the iron in the medium; alternatively, non-protein iron chelators such as 2,2Ϗ-bipyridine have been used to study siderophore cheating in *Burkholderia cenocepacia* [35]. Based on the outcome of our genome analysis, we also encourage authors investigating siderophore cheating to revise their sequencing data seeking for gene deletions that may responsible for important phenotypic changes, including the loss of exoprotease production and motility.

## Acknowledgements

R G-C research is funded by CONACYT grant CB 2017-2018 number A1-S-8530 and by PAPITT UNAM grant number IN214218, we are grateful with Cecilia Martinez Castillo for her technical assistance with some experiments.

## Notes

### Competing Interest Statement

The authors have declared no competing interest.

