## Supplementary material for "Iron limitation by transferrin promotes simultaneous cheating of pyoverdine and exoprotease in *Pseudomonas aeruginosa*": Table 1

| **Strain** | **Characteristics** | **Origin** |
| --- | --- | --- |
| PA14 | Wild-type strain, pyoverdine and exoprotease producer | [1] |
| PA14 *lasR* | PA14 with *lasR* interrupted by a transposon | [1] |
| CH1 | PA14 isolated from CAA TF, non-pyoverdine and non-exoprotease producer | This work |
| CH2 | PA14 isolated from CAA chelex, non-pyoverdine producer but exoprotease producer | This work |
| CH3 | PA14 isolated from CAA TF, pyoverdine producer and non-exoprotease producer | This work |
| WT1 | PA14 isolated from CAA TF, pyoverdine producer and exoprotease producer | This work |
| CH1-pUCP20 | CH1 complemented pUCP20 plasmid | This work |
| CH1-*lasR* | CH1 complemented with pUCP20-*lasR* | This work |
| CH3-pUCP20 | CH3 complemented pUCP20 plasmid | This work |
| CH3-*lasR* | CH3 complemented with pUCP20-*lasR* | This work |
| CH2-pUCP20 | CH2 complemented pUCP20 plasmid | This work |
| CH2-*pvdS* | CH3 complemented with pUCP20-*pvdS* | This work |
| PA01 | Wild-type strain, pyoverdine and exoprotease producer | [2] |

List of strains used in this work.

1.	Liberati NT, Urbach JM, Miyata S, Lee DG, Drenkard E, Wu G, et al. An ordered, nonredundant library of Pseudomonas aeruginosa strain PA14 transposon insertion mutants. *Proc Natl Acad Sci U S A* 2006.

2.	Chandler CE, Horspool AM, Hill PJ, Wozniak DJ, Schertzer JW, Rasko DA, et al. Genomic and phenotypic diversity among ten laboratory isolates of Pseudomonas aeruginosa PAO1. *J Bacteriol* 2019.
