## supplementary figures for "Iron limitation by transferrin promotes simultaneous cheating of pyoverdine and exoprotease in *Pseudomonas aeruginosa*"

### Slide 1
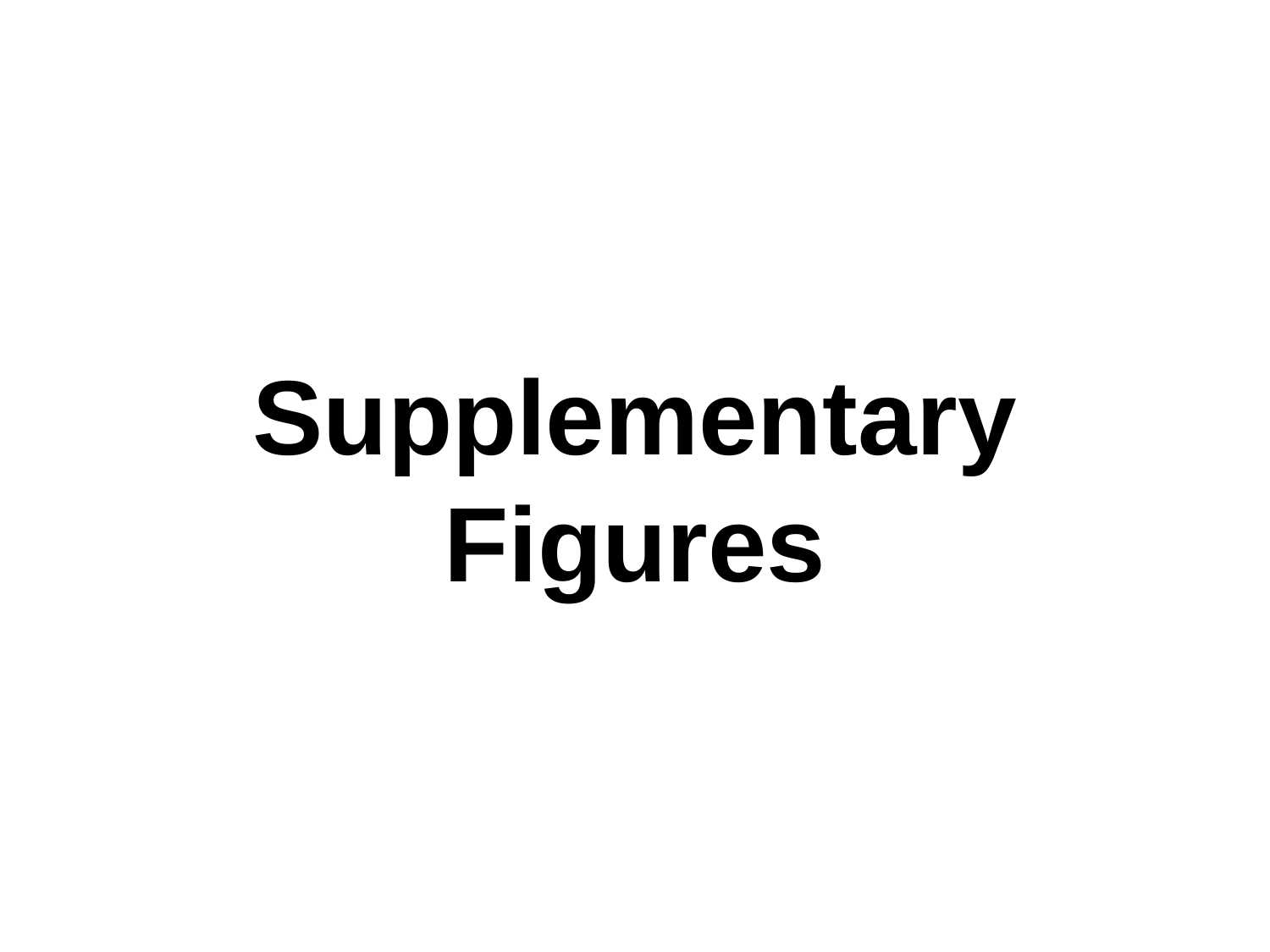

#### Supplementary Figures

### Slide 2
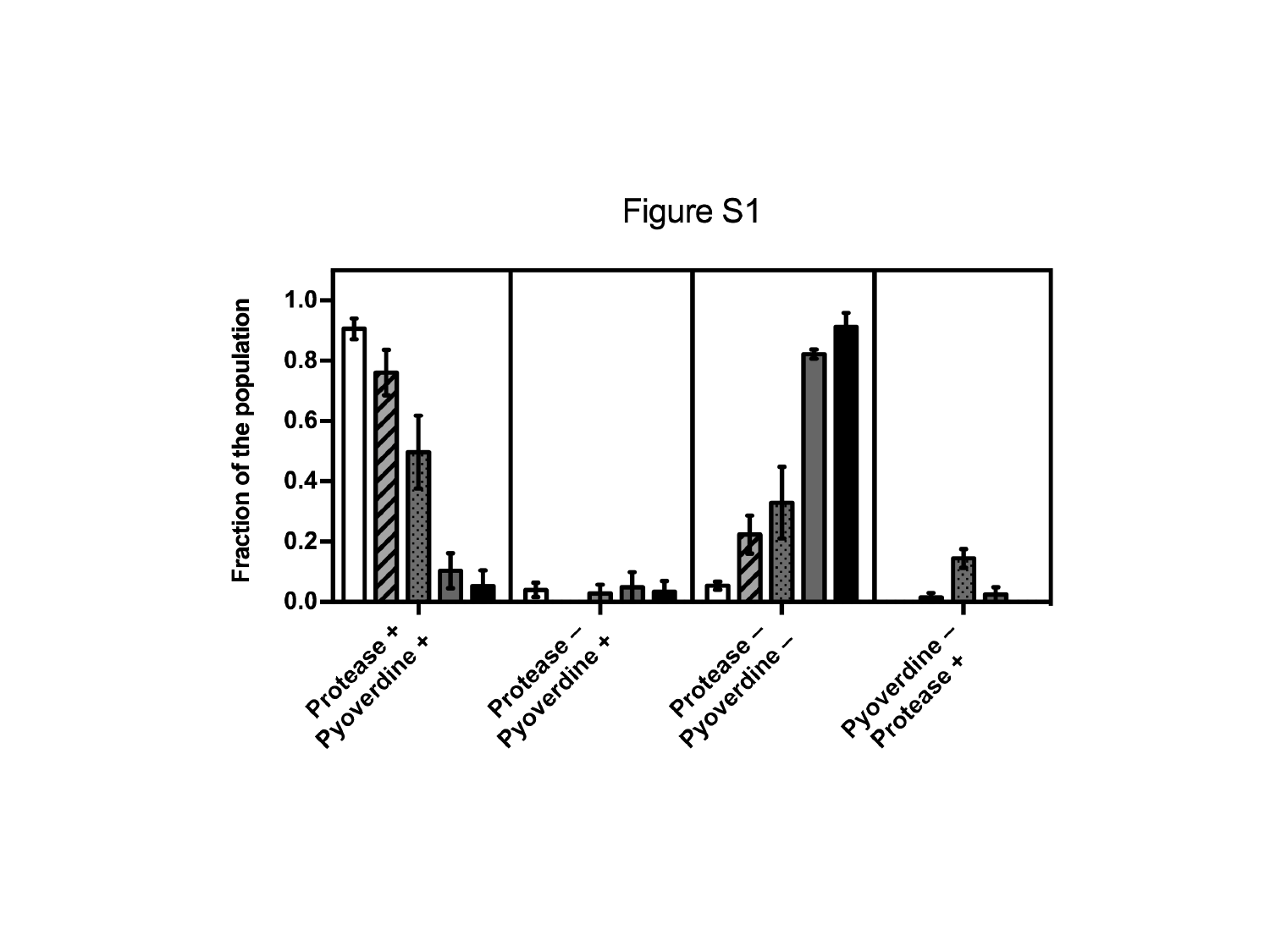

### Slide 3
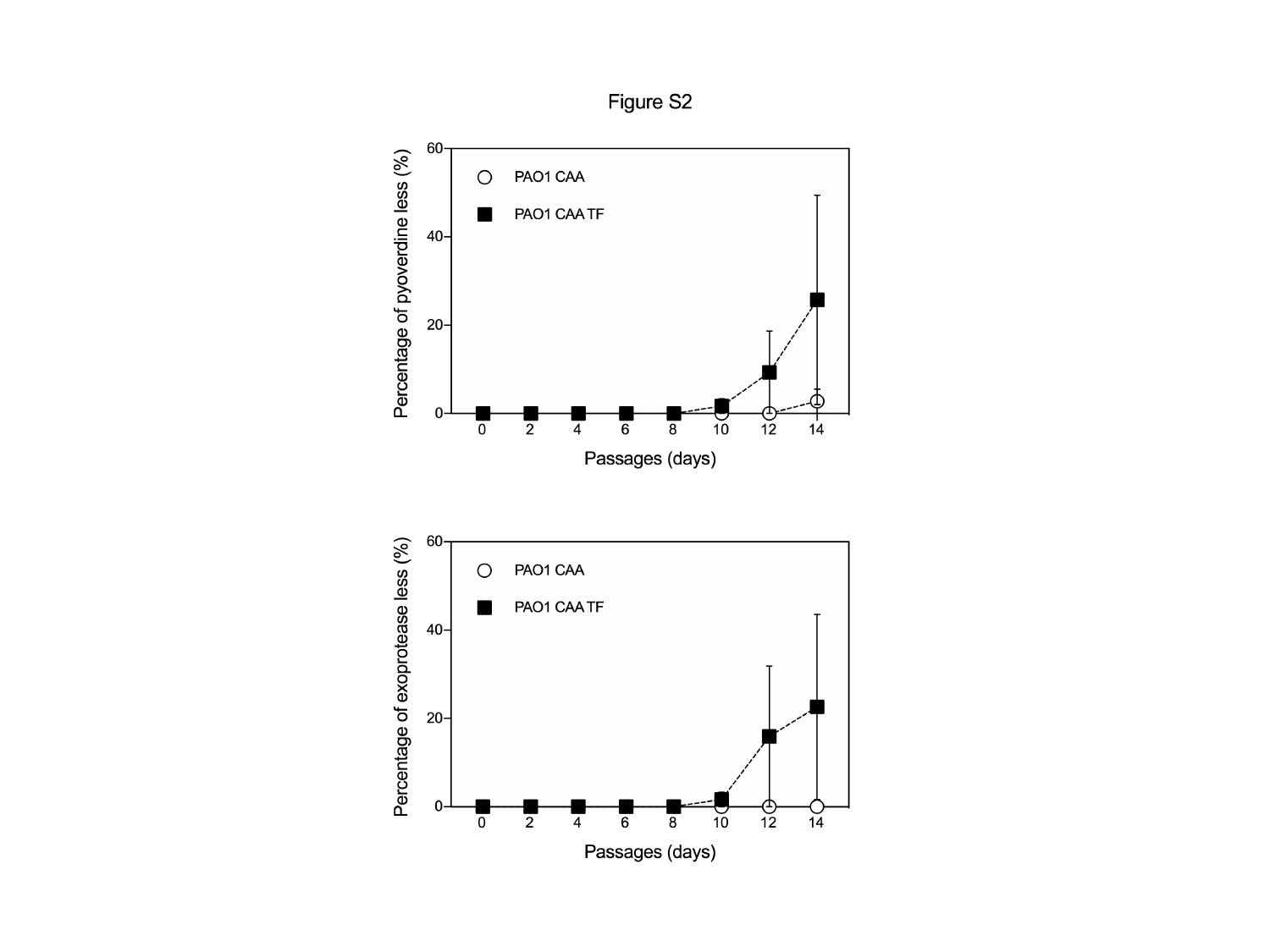

### Slide 4
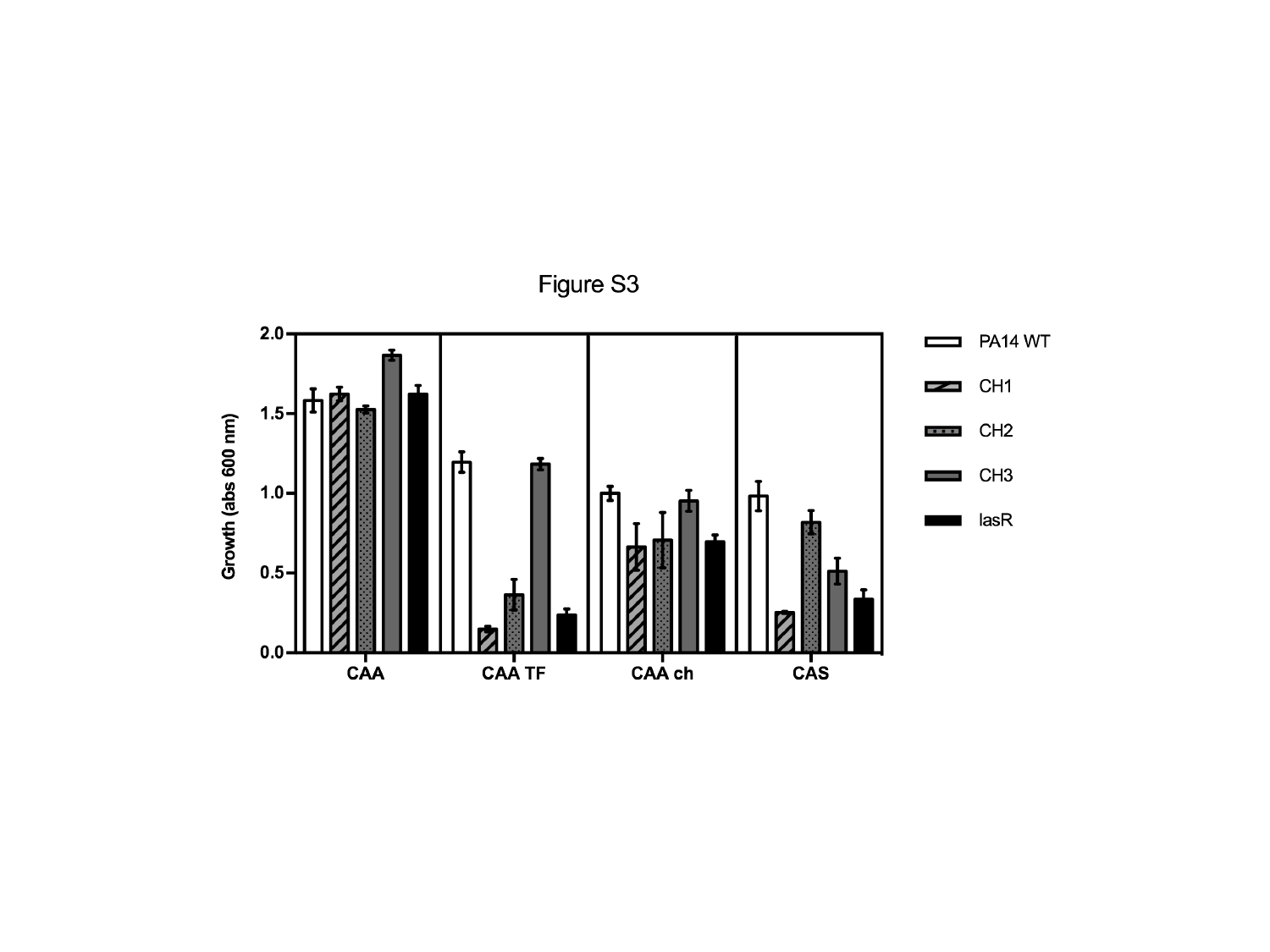

### Slide 5
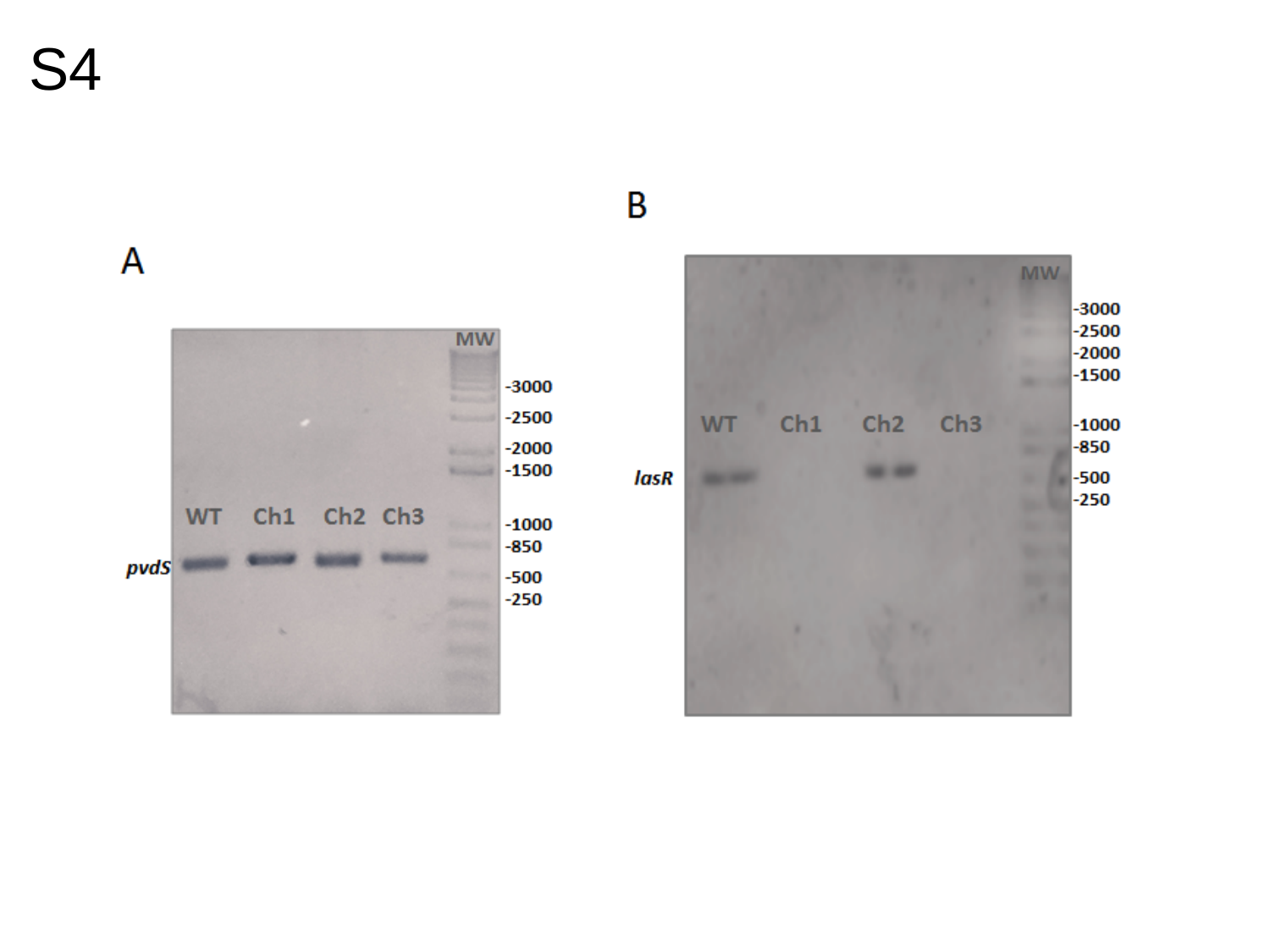

S4

### Slide 6
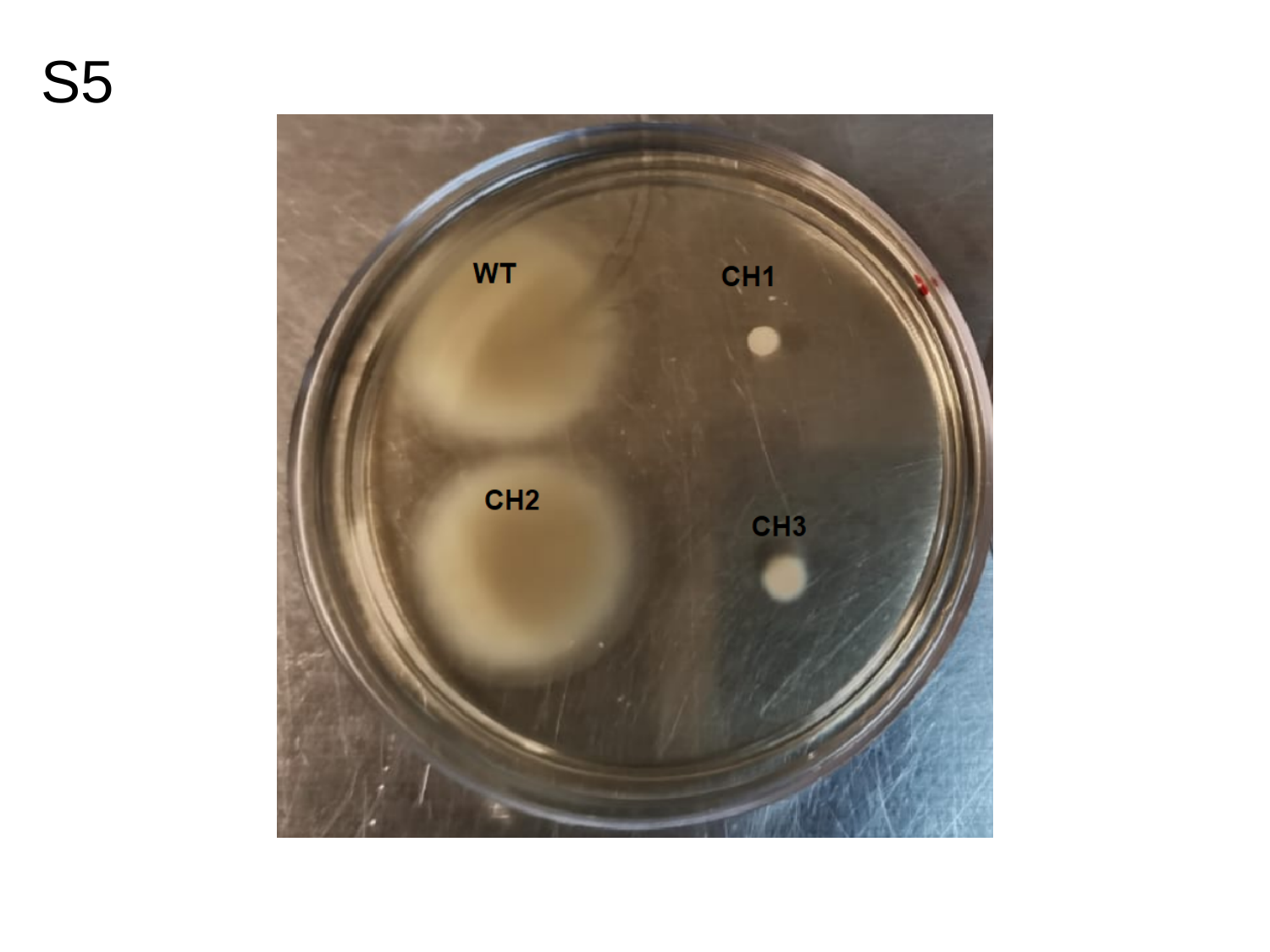

S5

### Slide 7
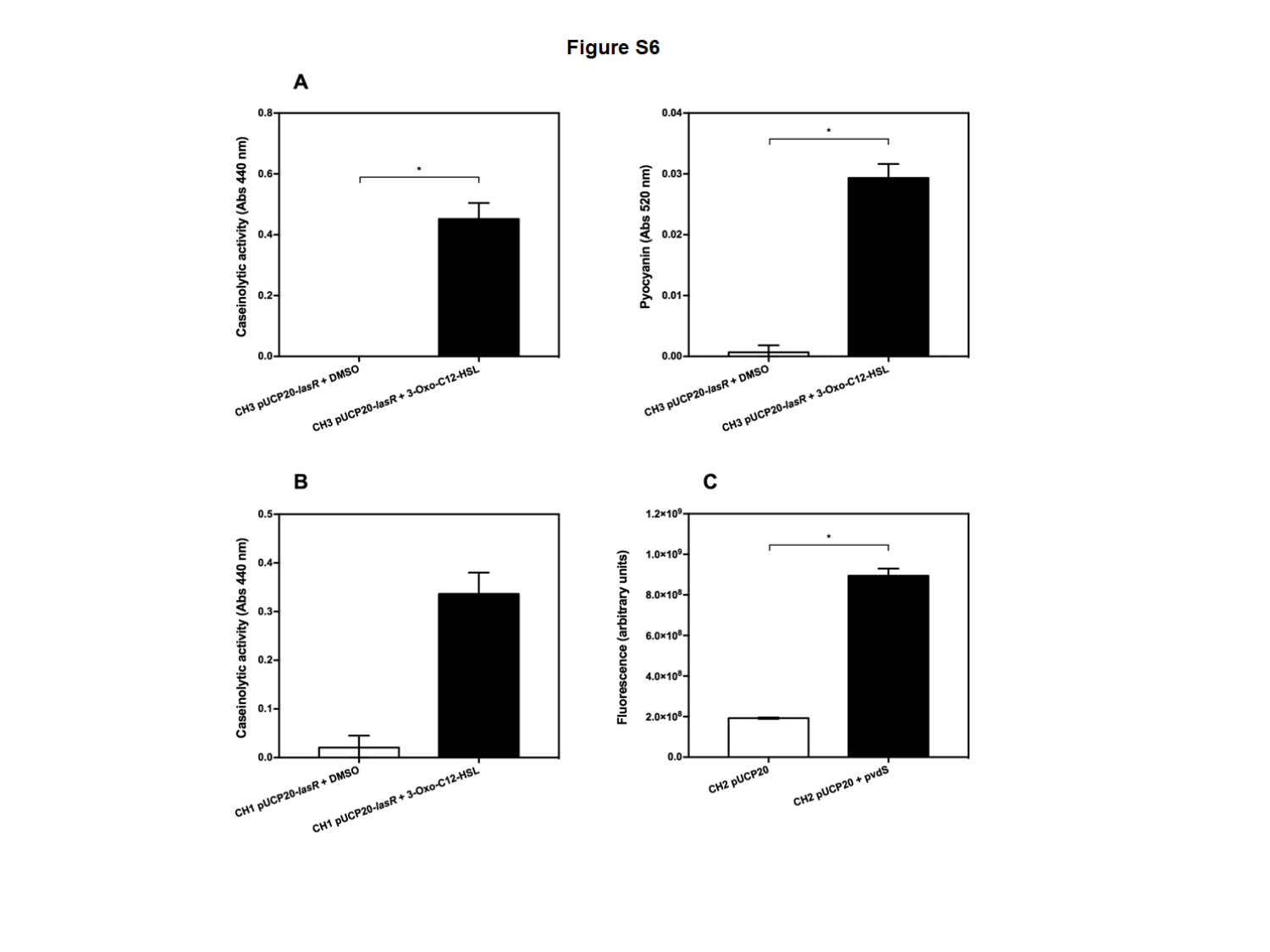

### Slide 8
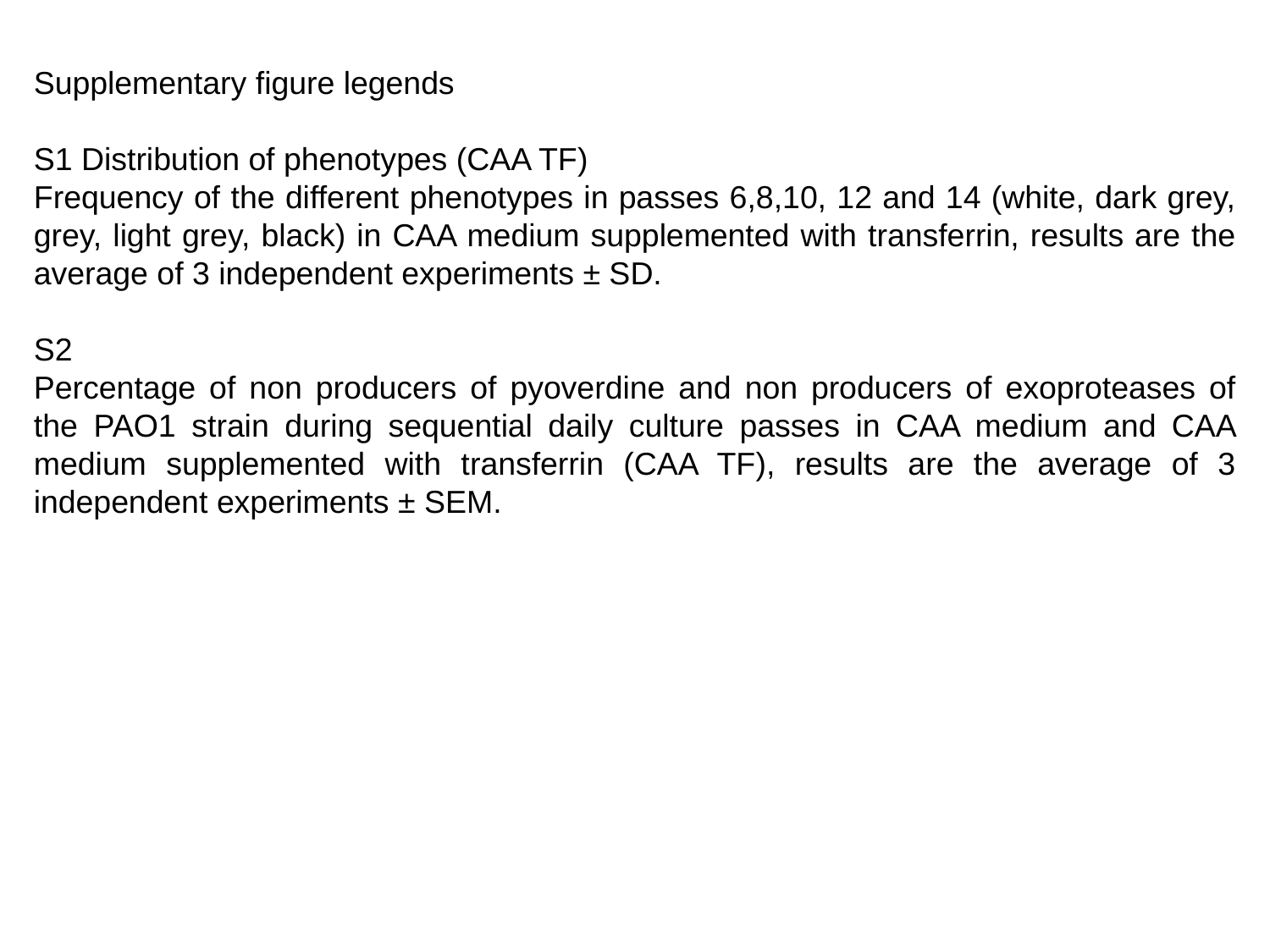

Supplementary figure legends
S1 Distribution of phenotypes (CAA TF)
Frequency of the different phenotypes in passes 6,8,10, 12 and 14 (white, dark grey, grey, light grey, black) in CAA medium supplemented with transferrin, results are the average of 3 independent experiments ± SD.
S2
Percentage of non producers of pyoverdine and non producers of exoproteases of the PAO1 strain during sequential daily culture passes in CAA medium and CAA medium supplemented with transferrin (CAA TF), results are the average of 3 independent experiments ± SEM.

### Slide 9
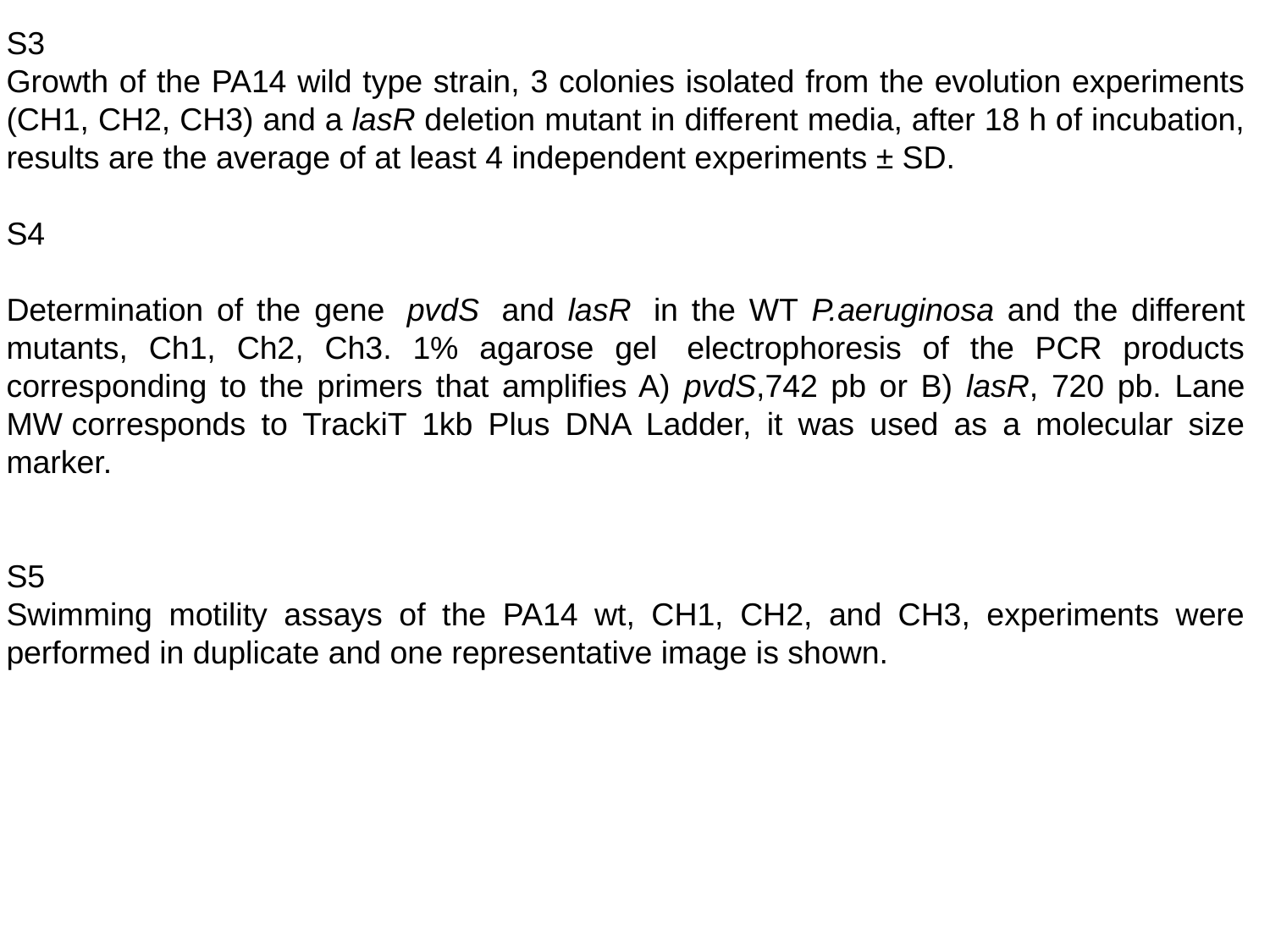

S3
Growth of the PA14 wild type strain, 3 colonies isolated from the evolution experiments (CH1, CH2, CH3) and a lasR deletion mutant in different media, after 18 h of incubation, results are the average of at least 4 independent experiments ± SD.
S4
Determination of the gene  pvdS  and lasR  in the WT P.aeruginosa and the different mutants, Ch1, Ch2, Ch3. 1% agarose gel  electrophoresis of the PCR products corresponding to the primers that amplifies A) pvdS,742 pb or B) lasR, 720 pb. Lane MW corresponds to TrackiT 1kb Plus DNA Ladder, it was used as a molecular size marker.
S5
Swimming motility assays of the PA14 wt, CH1, CH2, and CH3, experiments were performed in duplicate and one representative image is shown.

### Slide 10
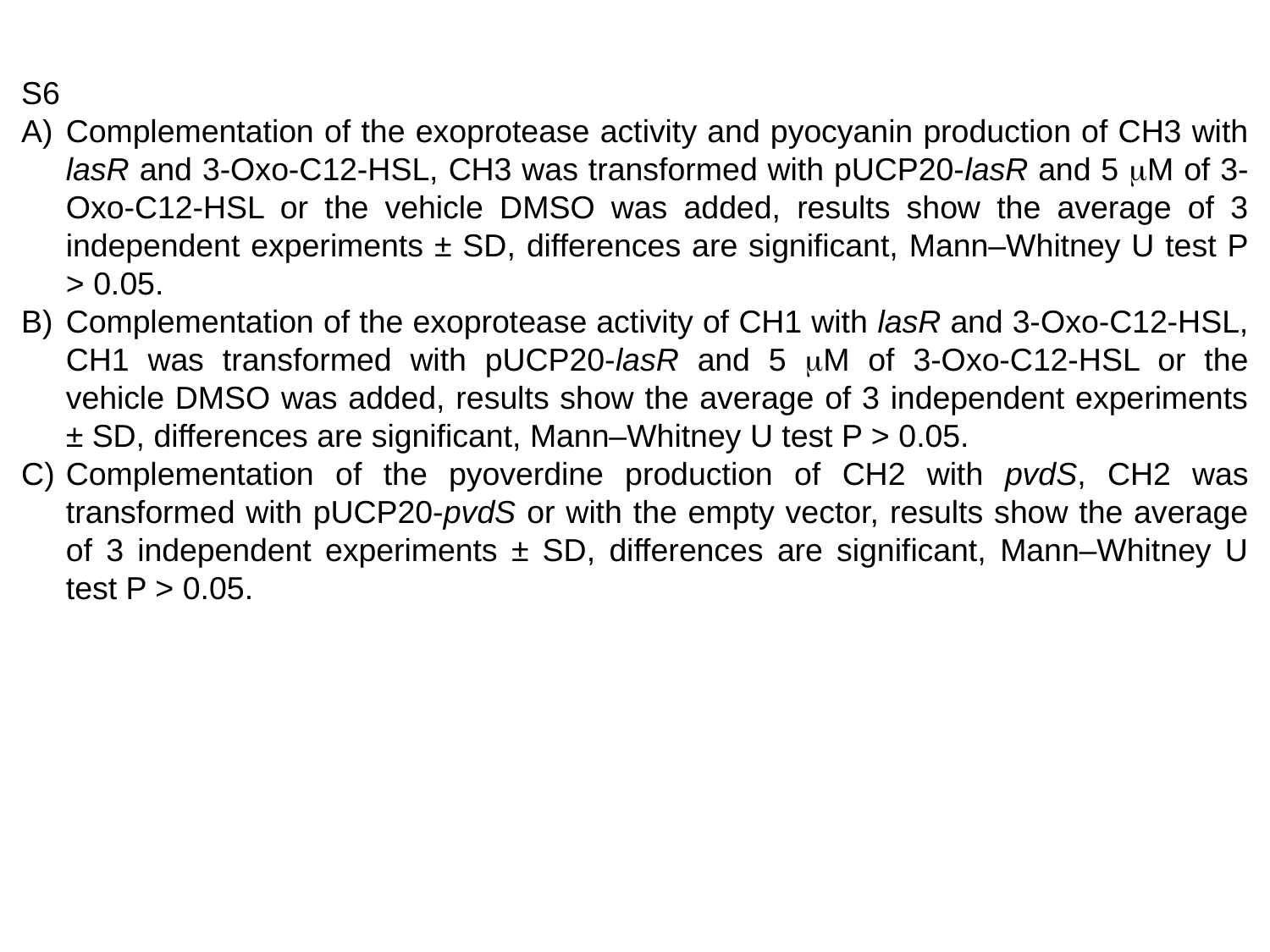

S6
Complementation of the exoprotease activity and pyocyanin production of CH3 with lasR and 3-Oxo-C12-HSL, CH3 was transformed with pUCP20-lasR and 5 μM of 3-Oxo-C12-HSL or the vehicle DMSO was added, results show the average of 3 independent experiments ± SD, differences are significant, Mann–Whitney U test P > 0.05.
Complementation of the exoprotease activity of CH1 with lasR and 3-Oxo-C12-HSL, CH1 was transformed with pUCP20-lasR and 5 μM of 3-Oxo-C12-HSL or the vehicle DMSO was added, results show the average of 3 independent experiments ± SD, differences are significant, Mann–Whitney U test P > 0.05.
Complementation of the pyoverdine production of CH2 with pvdS, CH2 was transformed with pUCP20-pvdS or with the empty vector, results show the average of 3 independent experiments ± SD, differences are significant, Mann–Whitney U test P > 0.05.
